# APP functions at the endomembrane system

**DOI:** 10.64898/2026.11.01.763500

**Authors:** Zichen Wang, Zhenli Fu, Guoqing Lai, Qiwen Zheng, Wenliangyu Li, Zhihan Lei, Xinlin Xia, Xuehan Zhu, Ying Lou, Derong Liu, Xiubo Du, Qiong Liu, Binbin Zheng, Nan Li

**Affiliations:** Shenzhen Key Laboratory of Marine Biotechnology and Ecology, Brain Disease and Big Data Research Institute, College of Life Sciences and Oceanography, Shenzhen University, Shenzhen 518055, China; Instrumental Analysis Center of Shenzhen University, Shenzhen University, Shenzhen 518055, China; Shenzhen Bay Laboratory, Shenzhen 518036, China

**Keywords:** APP, FE65, ATF4, Golgi apparatus, Endoplasmic reticulum stress, autophagy, APP-CTFs

## Abstract

**INTRODUCTION:** APP is a type I transmembrane protein, and its intracellular domain (AICD) interacts with FE65, X11, and mDAB1, et al., through the YENPTY motif. Accordingly, it has been proposed that the functions of APP are similar to Notch, which regulates gene expression through releasing NICD after combining with its ligands. However, the detailed underlying mechanisms of how FE65 is regulated by APP, as well as their target genes, remain unknown.

**METHODS:** We overexpressed APP695 in N2a cells, and deleted/mutated APP in H4 cells by CRISPR-Cas9, respectively. The impacts of APP overexpression and ablation were detected by proteome comparison and immunoblotting. Thereafter, the subcellular localizations of APP and its binding protein FE65 were examined by immunofluorescence.

**RESULTS:** APP overexpression not only resulted in its retention in Golgi apparatus (GA) and Endoplasmic reticulum (ER), but also induced FE65 clustering. In addition, it significantly reduced the protein level of ATF4 in N2a cells. On the contrary, knockdown of FE65 restored the protein level of ATF4. More importantly, APP overexpression also impaired autophagy. In contrast, knockout of APP rescued LC3 conversion. Of note, the APP Swedish mutant did not impact ER stress or the autophagic flux. Nevertheless, it facilitated the cleavage of APP-CTFs by γ-secretase.

**DISCUSSION:** The evidence presented in the current study demonstrates that APP is involved in stabilizing the endomembrane system and the subcellular transport of FE65. More importantly, the role of APP in regulating autophagy indicated that full-length APP could be considered a therapeutic target for preventing the development of AD.

## 1 Background

Amyloid-precursor protein (APP) is a type I transmembrane protein, which is encoded by the *APP* gene that is located on chromosome 21. Many mutations of *APP* are identified as autosomal dominant genetic risk factors for familial Alzheimer’s disease (FAD) and/or cerebral amyloid angiopathy (CAA) because they tend to result in the overload of amyloid-beta (Aβ) ^1^. According to the alternative splicing of *APP* mRNA, there are three isoforms of APP mainly expressed in the brain, which contain 770, 751, and 695 amino acids, respectively. Compared with APP770, APP751 misses the Ox-2 domain, APP695 lacks the Ox-2 antigen domain and Kunitz protease inhibitor (KPI) domain in the extracellular compartment. However, they share the same transmembrane domain and C-terminal fragment (CTF). All of them can be hydrolyzed by α-secretase to generate the soluble APPα fragment and the C-terminal fragment containing 83 a.a. (CTF83), the latter one can be further hydrolyzed by γ-secretase to produce a P3 fragment, and an APP intracellular domain (AICD) ^2^. Alternatively, they can also be hydrolyzed by β-secretase to generate the soluble APPβ fragment and the C-terminal fragment containing 99 a.a. (CTF99), then the latter one can be further hydrolyzed by γ-secretase to produce the Aβ and AICD ^3^. Of note, there is another cleavage site of β-secretase, which produces the CTF89 ^4^. In addition to α, β, and γ−secretase, APP was also found to be the substrate of caspase-3 ^5–8^, matrix metalloproteinase (MMP) ^9,10^, HtrA2 ^11^, RHBDL4 ^12^, Meprin-β ^13^, as well as asparagine endopeptidase (AEP) ^14^. However, whether the proteolytic processes are involved in generating unique fragments of APP with specific biological activity or degradation of APP is not fully understood so far.

In terms of the hydrolyzing process of APP by α− and γ−secretase, it showed great similarity with Notch, which is proteolyzed by ADAM10 and γ−secretase to generate a Notch intracellular domain (NICD) when Notch binds to its ligands, such as Delta or Serrate/Jagged families. Subsequently, NICD exhibits transcriptional activity by forming a complex with DNA-binding protein CBF1, Suppressor of Hairless, Lag-1 (CSL) and the coactivator Mastermind (MAM) to regulate the transcription of target genes ^15^. Indeed, the interaction of APP intracellular domain (AICD) with FE65 was first reported in 1995 ^16^. Thereafter, the transcriptional activity of AICD was identified by Cao and Südhof in 2001 through co-transfecting Gal4-AICD into cell lines that contained the constitutive β-galactosidase expression plasmid. It was shown that FE65 stimulated transcription of β-galactosidase together with Gal4-AICD and histone acetyltransferase Tip60 ^17^. However, the mechanism underlying FE65 activation by AICD was quite controversial. For example, Cao and Südhof demonstrated that the nuclear translocation of AICD may be dispensable for the transcriptional activity of FE65. Instead, the membrane-tethered AICD is essential for FE65 activation ^18^. Consistently, Hass and Yankner also reported that AICD did not activate transcription, whereas APP recruited Tip60 to the membrane compartment to induce the formation of FE65-Tip60 complex in a CDK5-dependent manner ^19^. Interestingly, the phosphorylation of APP at Thr-668 was found to impact the interaction of APP with FE65 by causing a conformational change in the cytoplasmic domain that contains the YENPTY motif ^20^, thus resulting in liberation of membrane-bound FE65, which was then translocated into the nucleus ^21^.

Revealing the biological function of APP may contribute to the understanding of the pathology of AD and CAA. Therefore, we analyzed the impacts of APP overexpression on N2a cells and the influences of APP knockout on H4 cells, respectively. Interestingly, the observations in the current study demonstrated that APP physiologically interacts with FE65, and further manifested that the interaction of APP with FE65 is involved in regulating the homeostasis of the endomembrane system.

## 2 Methods

### 2.1 Chemicals and reagents

Dulbecco’s modified Eagle medium (DMEM), Opti-MEM I Reduced Serum Medium, and Geneticin selective antibiotic (G418 sulfate) were purchased from Gibco (Thermo Fisher Scientific, Waltham, MA, USA). Trypsin-EDTA (0.25%) was obtained from Biosharp (Hefei, China). Dimethyl sulfoxide (DMSO) was purchased from Solarbio (Beijing, China). Penicillin-streptomycin solution was obtained from Vivacell (Shanghai, China). Fetal bovine serum (FBS) was purchased from ExCell Bio (Shanghai, China). Control siRNA-A and mouse FE65 siRNA were purchased from Santa Cruz Biotechnology (Dallas, TX, USA). The Pierce BCA Protein Assay Kit was obtained from Thermo Fisher Scientific (Waltham, MA, USA). Alexa Fluor 555-conjugated AffiniPure goat anti-mouse IgG (H+L), Alexa Fluor 555-conjugated AffiniPure goat anti-rabbit IgG (H+L), Alexa Fluor 488-conjugated AffiniPure goat anti-rabbit IgG (H+L), and Alexa Fluor 488-conjugated AffiniPure goat anti-mouse IgG (H+L) were purchased from Jackson ImmunoResearch (West Grove, PA, USA).Source data for all primary antibodies employed in Western blot and immunofluorescence assays, including supplier details and catalog numbers, are compiled in Table 1.

### 2.2 Cell culture

The mouse neuroblastoma Neuro-2a (N2a) cell line was cultured in a 1:1 mixture of DMEM and Opti-MEM I supplemented with 6% (v/ v) fetal bovine serum (FBS) and 1% (v/v) penicillin-streptomycin (Gibco) at 37 ℃ in a humidified incubator with 5% CO_2_. N2a cells stably transfected with human APP Swedish mutant (695 amino acids; referred to as N2a-APPswe or N2a-SW) were maintained under identical conditions, except that 400 μg/mL G418 was added to the medium for continuous selection. Cells were passaged or harvested for protein extraction when they reached approximately 80% confluence. N2a cell lines transfected with empty vector pCB6 (N2a/vector), or APP Swedish mutant (N2a/APPswe) were kindly gifted by Prof. Huaxi Xu (Sanford Burnham Prebys Medical Discovery Institute, Xiamen University) ^22^

### 2.3 Knockdown and overexpression of FE65

For FE65 knockdown, cells were transfected with 50 nM control siRNA-A or mouse FE65 siRNA using Lipofectamine LTX PLUS reagent (Invitrogen, A1260) according to the manufacturer’s instructions. siRNA-reagent complexes were prepared in Opti-MEM I reduced serum medium and incubated for 5 min at room temperature before being added to the two types of cells. After 8 h transfection, the medium was replaced with complete growth medium, and cells were incubated for an additional 18 h before sample collection. For FE65-EGFP fusion protein overexpression, the empty plasmid or FE65-EGFP containing plasmid at 10 µg per dish were transfected by Lipofectamine 3000 (Invitrogen, L3000015). 24h after, the medium was replaced with PBS, the subcellular localization of FE65 was examined by confocal laser scanning microscopy.

### 2.4 Protein lysis extraction

Cells were washed twice with ice-cold PBS and lysed by sonication on ice (8 s twice with a 1 min interval) in radioimmunoprecipitation assay (RIPA) lysis solution containing 1 mM phenylmethylsulphonyl fluoride (PMSF; Thermo Fisher Scientific, Waltham, MA, USA) supplemented with phosphatase and proteinase inhibitors. Homogenates were centrifuged at 12,000 × g for 30 min at 4 °C, thereafter the supernatants were collected for subsequent analysis. And the samples were collected and stored at −80 °C.

### 2.5 Immunoblot analysis

Protein concentrations were determined using the Pierce BCA Protein Assay Kit (Thermo Fisher Scientific, catalog no. 23225). Equal amounts of protein (20 μg per lane) were separated by 10% sodium dodecyl sulfate-polyacrylamide gel electrophoresis (SDS-PAGE) and electro-transferred onto polyvinylidene difluoride (PVDF) membranes (Millipore, Burlington, MA, USA). Membranes were blocked with 5% (w/v) nonfat milk in Tris-buffered saline containing 0.1% Tween 20 (TBST) for 1 h at room temperature, followed by overnight incubation at 4 °C with primary antibodies diluted in primary antibody dilution buffer (Beyotime, catalog no. P0023A). After three 10 min washes with TBST, membranes were incubated with horseradish peroxidase (HRP)-conjugated secondary antibodies for 2 h at room temperature. Following being washed with TBST 10 min for four times, immunoreactive bands were visualized using an enhanced chemiluminescence (ECL) detection kit (Advansta, San Jose, CA, USA) and captured using a chemiluminescence imaging system. Band intensities were quantified by densitometric analysis using ImageJ software (National Institutes of Health, Bethesda, MD, USA) and normalized to the corresponding loading controls (β-actin or GAPDH).

### 2.6 Immunofluorescence

Cells were seeded onto 20 mm tissue culture-treated glass-bottom dishes and allowed to grow to approximately 60% confluence. Cells were washed three times with PBS, fixed with 4% paraformaldehyde (PFA) in PBS for 15 min at room temperature, permeabilized with 0.2% (v/v) Triton X-100 in PBS for 10 min, and blocked with 10% (v/v) normal goat serum in PBS for 1 h at room temperature. Cells were then incubated overnight at 4 °C with appropriately diluted primary antibodies.

After washing 5 min with PBS for three times, cells were incubated with Alexa Fluor 555-conjugated goat anti-mouse IgG (H+L) or Alexa Fluor 488-conjugated goat anti-rabbit IgG (H+L) secondary antibodies for 1 h at room temperature in the dark. Then, following being washed 5 min with PBS for three times, nuclei were counterstained with DAPI (1 μg/ mL) for 10 min. After three times final washes with PBS, 5 min each time, cells were mounted with antifade mounting medium and imaged using a Zeiss Elyra 7 Lattice SIM with LSM 980 confocal microscope (Carl Zeiss, Oberkochen, Germany).

### 2.7 CRISPR editing and single-cell sorting of H4-APPswe and H4-APPKO cell lines

H4 human neuroglioma cells were seeded in 48-well plates at a density of 4 × 10⁴ cells per well in complete growth medium and allowed to adhere overnight at 37 °C in a humidified atmosphere containing 5% CO₂. Genome editing was performed using Lipofectamine CRISPRMAX Cas9 Transfection Reagent (Thermo Fisher Scientific) according to the manufacturer’s instructions, with reagent volumes scaled to the 48-well format. Recombinant Streptococcus pyogenes Cas9 nuclease (GenScript) and an APP-targeting single-guide RNA (sgAPP), both at 10 μM, were incubated together at room temperature for 10 min to form Cas9–sgRNA ribonucleoprotein (RNP) complexes. The spacer sequence of sgAPP is provided in Table 2.

To generate APP-knockout (APP-KO) cells, Cas9–sgRNA RNP complexes were transfected without a repair template. To generate the Swedish mutation knock-in line, the RNP complexes were co-transfected with 0.5 μg per well of a single-stranded oligodeoxynucleotide (ssODN) homology-directed repair donor harboring the Swedish double mutation at residues 670 and 671 (KM670/671NL). The donor sequence with corresponding homology arms is provided in Table 1. At 12 h after transfection, the medium was replaced with fresh complete growth medium, and the cells were cultured for an additional 36 h.

At 48 h after transfection, the cells were washed with PBS and dissociated into a single-cell suspension using trypsin–EDTA. After neutralization with complete growth medium, the cell suspension was passed through a 40-μm cell strainer. Singlet cells were identified by sequential gating based on forward- and side-scatter characteristics, followed by doublet exclusion, and sorted using a FACSAria III cell sorter (BD Biosciences). Single cells were deposited individually into 96-well plates, allowed to undergo clonal expansion and subsequently transferred to larger culture vessels.

Genomic DNA was isolated from a portion of each expanded clone, and the genomic region encompassing the Cas9 cleavage site was amplified by PCR using the primers listed in Table 3. The resulting PCR products were sequenced to determine the genotype of each clone. APP-KO clones were identified by frameshifting insertions or deletions affecting all detectable APP alleles, whereas APPswe clones were identified by the presence of the intended KM670/671NL substitutions at the targeted locus.

For DAPT treatment, DAPT was dissolved in DMSO to prepare a stock solution and diluted to the working concentration in complete growth medium routinely used for H4 cells immediately before use. H4 cells were treated with 100 nM DAPT for 4 h, cells in the vehicle group were incubated in medium containing an equal volume of DMSO.

### 2.8 Proteome analysis

Enzymatic Digestion: Transfer 50 μg of protein sample into a new EP tube. Add 100 μL of denaturant (8 M urea), then add 10 μL of 100 mM DTT. Reduce protein at 37°C for 2 h (denaturation time and temperature depend on the denaturant used). Add 10 μL of 500 mM IAA in the dark for 15 min. Thereafter, add another 10 μL of 100 mM DTT, and react at room temperature for 15 min. Add pre-washed magnetic beads (25 μL–500 μg, bead-to-protein ratio 1:10), followed by an equal volume of acetonitrile. Vortex at room temperature for 30 min (1200 rpm). Centrifuge for 2 seconds, place on a magnetic stand for 2 min, and transfer the supernatant to a new EP tube. Add 180 μL of 80% acetonitrile to wash, and vortex at room temperature for 1 min. Centrifuge for 2 seconds, place on a magnetic stand for 2 min, and transfer the supernatant to a new EP tube (repeat twice). Place the tube in a fume hood to air-dry for 5 min. Add 1 μg of trypsin (in 200 μL of 25 mM ammonium bicarbonate). Mix by pipetting, sonicate for 5 min, and digest overnight at 37°C with shaking (600 rpm). Centrifuge at 20,000 rpm for 2 min, place on a magnetic stand for 2 min, and transfer the supernatant to a new EP tube. Add 200 μL of 2% DMSO to further elute the peptide sample. Sonicate for 5 min and vortex for 5 min. Centrifuge for 2 seconds, place on a magnetic stand for 2 min, and transfer the supernatant to a new EP tube. Combine the two eluates, dry by rotary evaporation, and proceed to mass spectrometry analysis.

Liquid chromatography-mass spectrometry: Reconstitute the dried peptide sample in a 0.1% formic acid-water solution. Liquid chromatography separation was performed using a Vanquish Neo nano-HPLC system. Mobile phase A consisted of 0.1% formic acid in water, and mobile phase B consisted of 0.1% formic acid in acetonitrile. The trap column (300 µm × 0.5 cm, RP-C18, Thermo Fisher Scientific) was equilibrated with 100% mobile phase A.

Samples were loaded via an autosampler and trapped onto the trap column, followed by analytical separation on a PepMap column (2 µm, 150 µm × 15 cm, Thermo Fisher Scientific) at a flow rate of 2.5 µL/min. The gradient elution conditions were as follows: 0.0–4.2 min, 4–25% B; 4.2–5.8 min, 25–35% B; 5.8–6.2 min, 35–99% B; and 6.2–6.9 min, 99% B. Following separation, the enzymatic digests were analyzed on an Orbitrap Astral Zoom mass spectrometer (Thermo Fisher Scientific, USA) operating in Data-Independent Acquisition (DIA) mode. Full-scan resolution was set to 240,000 over m/z range of 380–980. The Automatic Gain Control (AGC) target value was set to 500%. The maximum injection time (maxIT) was set to 2 ms. A total of 300 isolation windows with a width of 2 m/z were utilized to cover the scan range from 380 to 980 m/z. Isolated precursor ions were fragmented via Higher-energy Collisional Dissociation (HCD) with a Normalized Collision Energy (NCE) of 25%.

Database search: Data processing and library-free search were performed using DIA-NN (version 2.3.2). In silico digestion was configured for trypsin specificity, allowing up to one missed cleavage. Carbamidomethylation of cysteine (C) was set as a static (fixed) modification, whereas protein N-terminal acetylation and methionine oxidation (M) were designated as dynamic (variable) modifications. Both MS1 and MS2 mass tolerances were automatically determined and optimized by DIA-NN during search execution.

### 2.9 Statistics

All statistical analyses were conducted using GraphPad Prism 8.0 software (GraphPad). Results are presented as means ± s.e.m. For comparison of two experimental conditions, Mann-Whitney two-tailed nonparametric test was performed. Kruskal-Wallis One-way ANOVA with post hoc test was performed for analysis of additional experimental groups. Two-way ANOVA with Post hoc test for multiple comparisons was performed to analysis the data of N2a and N2a SW cells that were treated with control siRNA and FE65 siRNA, respectively.

## 3 Results

### 3.1 Overexpression of APP increased its immature form in N2a cells

The APP695 containing Swedish mutation (K595N/M596L) was overexpressed in N2a cells (N2a SW) as previously reported ^23^. The protein level of APP was detected by immunoblot with an antibody that specifically recognizes the CTF of APP. Interestingly, the overexpressed APP was predominantly presented as an immature form compared with that in normal N2a cells (Figure 1A-C), indicating that it was less glycosylated in N2a SW cells ^24^.

**Figure 1.**
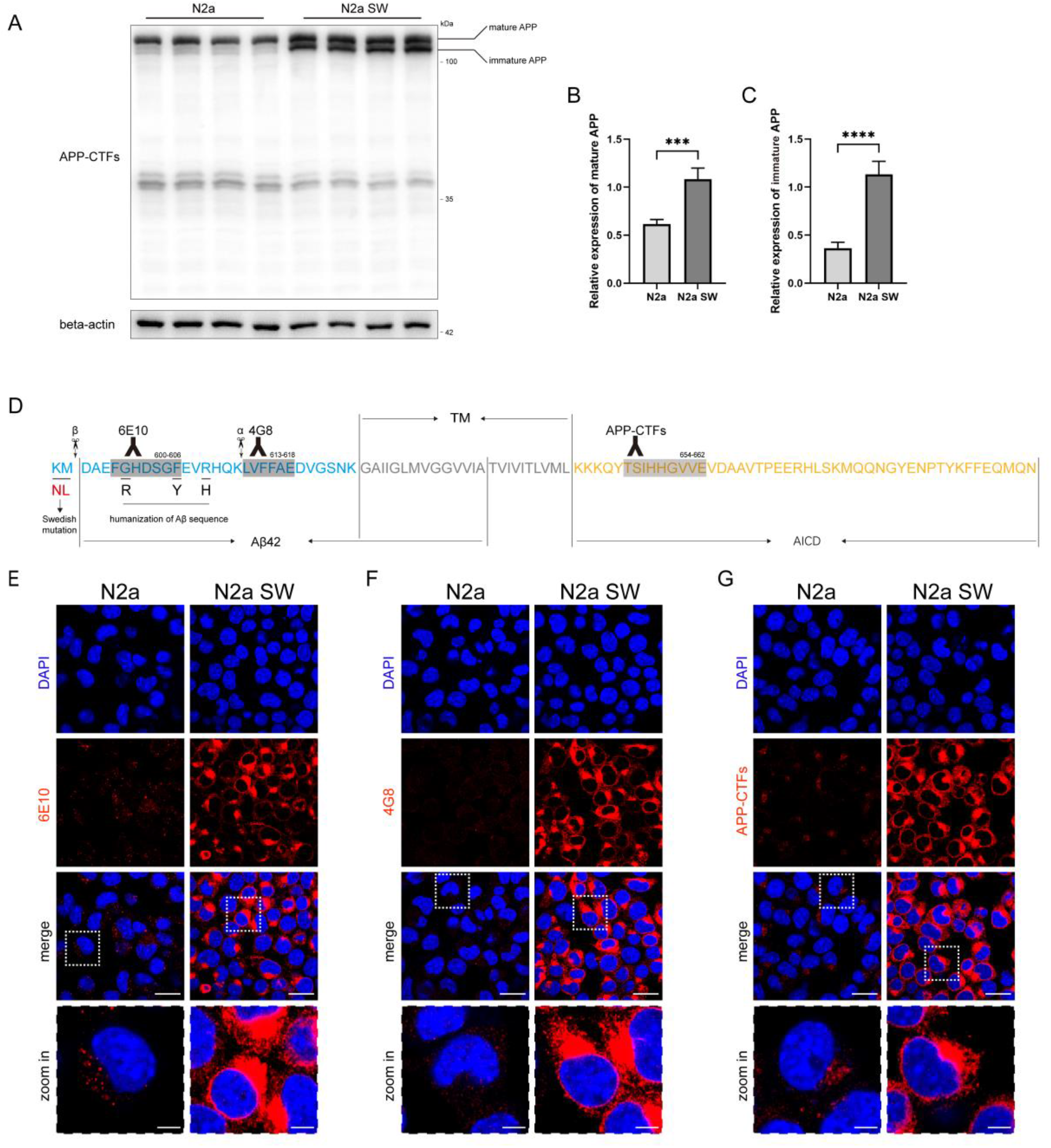
Overexpressed APP was retained around nuclei. **(A)** The protein levels of APP in the lysates from normal N2a cell and human APP695 overexpressing N2a cell (N2a SW) were examined by immunoblot using APP-CTFs specific antibody. **(B, C)** Quantification of mature APP/actin ratios and immature APP/actin ratios, respectively, (mean±SEM, *n*=4) (\*\*\**P*<0.001; \*\*\*\**P*<0.0001). **(D)** The schematic of APP epitopes that recognized by 6E10,4G8 and APP-CTFs antibodies (grey). The residues of Swedish mutation were denoted by red letters. The cleavage sites of α, β-secretases were indicated by scissors. The Aβ42 fragment, the transmembrane domain (TM), and the APP intracellular domain (AICD) in APP695 were demonstrated by arrows. **(E-G)** Representative images of immunofluorescence staining by 6E10 (E) 4G8 (F) and APP-CTFs (G) antibodies in N2a and N2a SW cells (red), the nuclei were visualized by DAPI (blue). Scale bars, 20 μm in original or 5 μm in zoomed in.

A previous study demonstrated that APP is transferred from the Golgi to the lysosome directly for degradation ^25^. The insufficient glycosylation of APP suggested that the transport of APP to the Golgi apparatus (GA) after its synthesis might be interrupted. Thus, we detected the subcellular localization of APP by 6E10, 4G8, and APP-CTF antibodies. 6E10 recognizes the N-terminus of APP-CTF99, 4G8 specifically binds to the N-terminus of APP-CTF83, whereas APP-CTF antibody combines with the AICD fragment of APP (Figure 1D). Interestingly, these antibodies showed different subcellular localization of APP fragments in N2a cells. The fluorescence signal of 6E10 antibody was mainly detected as puncta-like spots (Figure 1E left), which is similar to the feature of lysosome ^26^. The fluorescence signal of 4G8 in N2a cells was also detected in the cytosol, however, it was less distinct compared with that of 6E10 (Figure 1G left). In contrast, the fluorescence signal of APP-CTF antibody was detected as stacks adjacent to the nuclei (Figure 1G left), suggesting that full-length APP is mainly located on GA. However, in APP overexpressing N2a cells, these three antibodies consistently showed that APP was basically retained on GA (Figure 1E-G right). Strikingly, about half of these cells showed fluorescence signal around the nuclei, indicating that the immature APP was also partially retained on Endoplasmic reticulum (ER).

### 3.2 Overexpression of APP inhibited ER stress

To confirm the subcellular localization of APP, GA was visualized by GM130 specific antibody (Figure 2A, green). In N2a cells, the fluorescence signal of 6E10 (Figure 2A, red) was not colocalized with that of GM130 (Figure 2A, upper). In contrast, the fluorescence signal of these two antibodies showed extensive colocalization in N2a SW cells (Figure 2A, lower), illustrating that the overexpressed APP was mainly retained in GA. Of note, Structured Illumination Microscopy (SIM) showed that APP was mainly localized on the trans-Golgi, whereas GM130 was mainly distributed on the cis-Golgi **(Supplementary Figure 1A)**.

**Figure 2.**
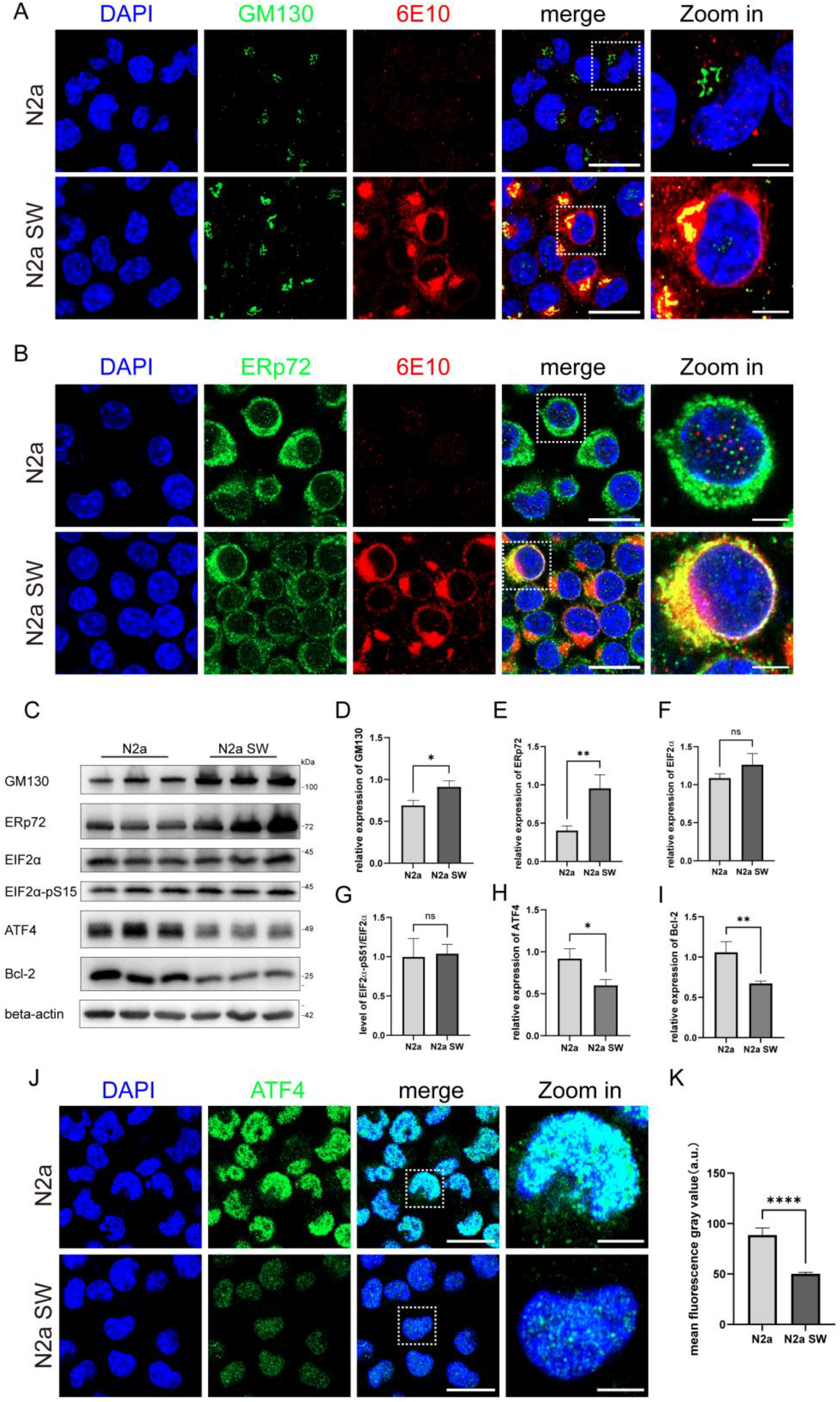
Retention of APP stabilized GA and ER. **(A)** Representative image of the immunofluorescence staining using GM130 (green) and 6E10 (red) antibodies in normal N2a cells (upper) and N2a SW cells (lower), respectively. The nuclei were visualized by DAPI (blue). Scale bars, 20 μm in original or 5 μm in zoomed in. **(B)** Representative image of the immunofluorescence staining using ERp72 (green) and 6E10 (red) antibodies in normal N2a cells (upper) and N2a SW cells (lower), respectively. The nuclei were visualized by DAPI (blue). Scale bars, 20 μm in original or 5 μm in zoomed in. **(C)** The protein levels of GM130, ERp72, eIF2α, S15 phosphorylated EIF2α, ATF4, Bcl-2 in the lysates from normal N2a cells and N2a SW cells were examined by immunoblot using specific antibodies, respectively. **(D-I)** Quantification of GM130/actin ratios (D), Erp72/actin ratios (E), EIF2α/actin ratios (F), EIF2α-pS15/EIF2α ratios (G), ATF4/actin ratios (H) and Bcl-2/actin ratios (I), respectively (mean±SEM, *n*=3) (\**P*<0.05; \*\**P*<0.01). **(J)** Representative image of immunofluorescence staining using ATF4 (green) antibody in normal N2a cells (upper) and N2a SW cells (lower), respectively. The nuclei were visualized by DAPI (blue). Scale bars, 20 μm in original or 5 μm in zoomed in. **(K)** Quantification of the ATF4 fluorescence intensity. (mean±SEM, *n*=8) (\*\*\*\**P*<0.0001)

Because overexpressed APP also distributed around nuclei in some N2a SW cells as mentioned above, we also stained ER by using ERp72 specific antibody (Figure 2B, green). In N2a cells, the fluorescence signal of 6E10 did not show colocalization with that of ERp72 (Figure 2B, upper). However, in N2a SW cells, the fluorescence signal of these two antibodies showed colocalization to some extent, especially in those that exhibited APP distribution around nuclei (Figure 2B lower).

Since the overexpressed APP was retained in GA, as well as in ER to some degree, we examined the protein level of GM130 and ERp72 by immunoblot (Figure 2C-E). The levels of these two proteins were significantly upregulated in N2a SW cells compared with normal N2a cells, indicating that overexpression of APP may contribute to stabilizing the structures of GA and ER. The morphology of GA in N2a SW cells became more compressed compared with N2a cells, which also corroborated this speculation **(Supplementary Figure 1B)**. These observations raised another question, i.e., whether overexpression of APP triggers ER stress. To address this question, we detected the phosphorylation level of eIF2α and protein level of ATF4, the well-known ER stress mediators, and their downstream protein Bcl-2 by immunoblot. Interestingly, although the results revealed that the phosphorylation level of eIF2α was not changed (Figure 2C, F, G), the protein levels of ATF4 and Bcl-2 were down-regulated in N2a SW cells compared with those in N2a cells (Figure 2C, H, I). Immunofluorescence of ATF4 also demonstrated that ATF4 was detected less in the nuclei of N2a SW cells than in the nuclei of N2a cells (Figure 2J, K).

### 3.3 Retention of APP promoted FE65 clustering

To understand the mechanism underlying the inhibition of ATF4 expression by overexpressed APP, we compared the proteome of N2a SW cells with that of N2a cells. Interestingly, besides APP, its interacting protein FE65 (Apbb1) was also significantly upregulated in N2a SW cells compared with N2a cells (Figure 3A, data are available at https://zenodo.org/records/23800114). It has been well established that the AICD domain of APP can promote the transcriptional activity of FE65, hence, we examined the protein level of FE65 by immunoblot. The result demonstrated that the level of FE65 was significantly increased in N2a SW cells compared with that of N2a cells (Figure 3B, C). Because APP was retained in GA and ER in N2a SW cells, we were curious about how it affected the subcellular localization of FE65. Therefore, we tracked the distribution of FE65 by immunofluorescence. Interestingly, the fluorescence signal of FE65 (Figure 3D, green) was visualized in the nucleus, cytosol, and protrusions of N2a cells (Figure 3D, upper), which is consistent with the finding that FE65 is associated with actin cytoskeleton regulators ^27^. However, in N2a SW cells it was clustered outside the nuclei and colocalized with the fluorescence of 6E10 (Figure 3D, lower) **(Video 1, 2)**. The intensity of the fluorescence signal was analyzed and presented as peaks in Figure 3E. Of note, N2a cells are derived from murine neurons, whether human FE65 could also be clustered by APP overexpression needs to be further examined. Thus, we overexpressed human FE65, which was linked with EGFP, in both N2a cells and N2a SW cells. The fluorescence of EGFP was detected in the nuclei of N2a cells, whereas in N2a SW cells it was detected outside the nuclei (Figure 3F). These observations further proved that the retention of APP in GA and ER promoted both murine and human FE65 clustering outside the nuclei, and also indicated that the protein level of FE65 played a critical role in its subcellular localization, i.e., under normal conditions, it was mainly localized at the boundary of the cells, with only a small portion distributed in nuclei. However, the majority of FE65 distributed in nuclei when it was overexpressed, unless overexpressed APP segregated it from entering the nuclei. Immunoblot by using FE65 specific antibody, which was also used in immunofluorescence staining, demonstrated that the FE65-EGFP fusion protein was efficiently overexpressed in both cell lines compared with cells transfected empty vector (Figure 3G).

**Figure 3.**
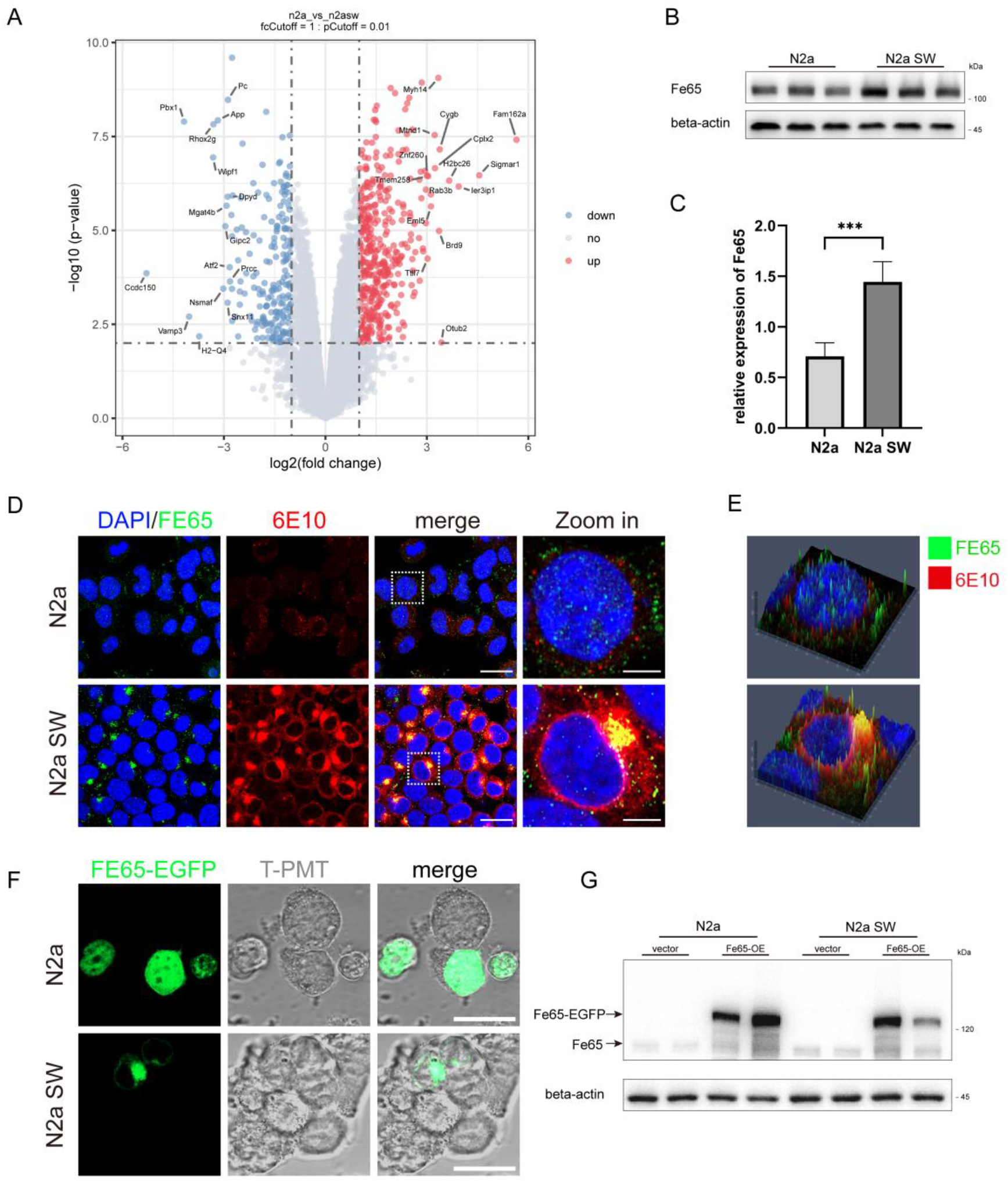
Overexpression of APP promoted FE65 clustering. **(A)** Volcano plots of the proteome of N2a cell compared with that of N2a SW cell (*n*=3, *P*<0.05, log^2^(Fold Change)>0.26). **(B)** The protein levels of FE65 in the lysates from normal N2a cells and N2a SW cells were examined by immunoblot using FE65 specific antibodies. **(C)** Quantification of FE65/actin ratios. (mean±SEM, *n*=3) (\*\*\**P*<0.001). **(D)** Representative image of the immunofluorescence staining using FE65 (green) and 6E10 (red) antibodies in normal N2a cells (upper) and N2a SW cells (lower), respectively. The nuclei were visualized by DAPI (blue). Scale bars, 20 μm in original or 5 μm in zoomed in. **(E)** Representative peak graph displayed the signal intensity profile of FE65 and 6E10 in a single cell of N2a and N2a SW, respectively. **(F)** Representative fluorescence images of FE65-EGFP fusion protein in N2a (upper) and N2a SW (lower) cells 24 h after FE65-EGFP plasmid was transfected. Scale bars, 10 μm **(G)** The protein level of FE65-EGFP and FE65 in the lysate of N2a and N2a SW that transfected with empty vector or FE65-EGFP for 24 h, respectively, were measured by immunoblot using FE65 specific antibody. Sample loading was normalized by beta-actin.

### 3.4 Knockdown of FE65 rescued protein level of ATF4 but not autophagy

It is of interest to learn how APP overexpression was related to inhibiting ATF4 expression. Since the interaction of FE65 and APP was found to be involved in regulating gene transcription as mentioned before, we attempted to knockdown FE65 by siRNA and examined the protein level of ATF4 again in N2a cells and N2a SW cells. Strikingly, accompanied by the down-regulation of FE65 by siRNA (Figure 4A, B), the protein level of ATF4 was significantly increased in both N2a cells and N2a SW cells (Figure 4A, C). The increased protein levels of CHOP and Bcl2 (Figure 4A, D, E), which are the well-known downstream effectors of ATF4 under ER stress, indicated that the upregulated ATF4 was functional.

**Figure 4.**
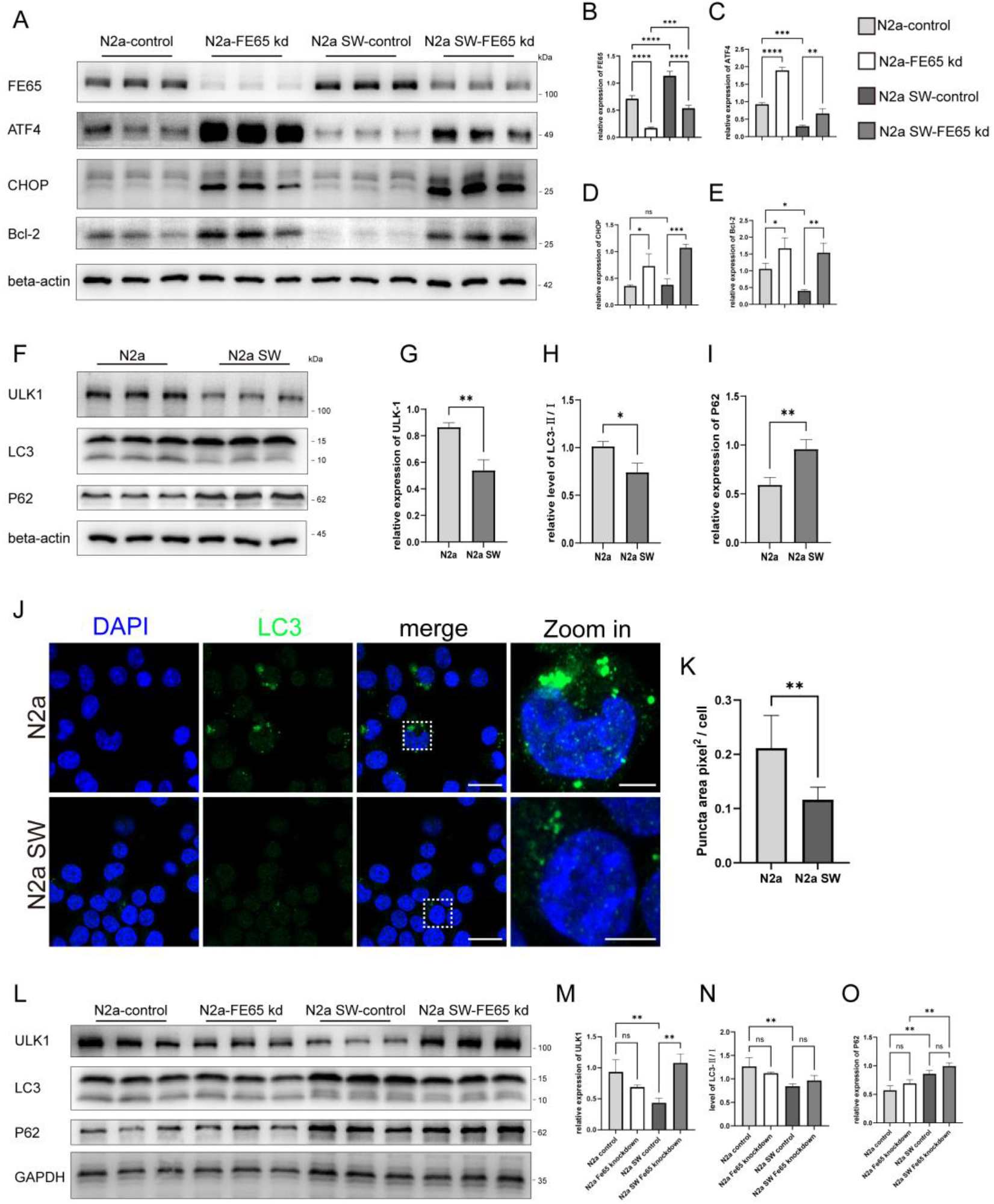
FE65 knockdown upregulated the protein level of ATF4. **(A)** The protein levels of FE65, ATF4, CHOP, Bcl-2 in the lysates from normal N2a cells and N2a SW cells that were transfected with random-siRNA or FE65-siRNA for 24 h, respectively, were examined by immunoblot using specific antibodies. **(B-E)** Quantification of FE65/actin ratios (B), ATF4/actin ratios (C), CHOP/actin ratios (D), Bcl-2/actin ratios (E), respectively, (mean±SEM, *n*=3) (\**P*<0.05; \*\**P*<0.01; \*\*\**P*<0.001; \*\*\*\**P*<0.0001). **(F)** The protein levels of ULK1, LC3, and P62 in the lysates from normal N2a cells and N2a SW cells were examined by immunoblot using specific antibodies. **(G-I)** Quantification of ULK1/actin ratios (G), LC3II/LC3I ratios (H), p62/actin ratios (I), respectively, (mean±SEM, *n*=3) (\**P*<0.05; \*\**P*<0.01). **(J)** Representative immunofluorescence staining using LC3 antibody (green) in N2A cells (upper) and N2a SW cells (lower). The nuclei were visualized by DAPI (blue). Scale bars, 20 μm in original or 5 μm in zoomed in. **(K)** Quantification of LC3 immunofluorescence puncta area pixel^2^ per cell, (mean±SEM, *n*=8) (\*\**P*<0.01). **(L)** The protein levels of ULK1, LC3, and P62 in the lysates from normal N2a cells and N2a SW cells that were transfected with random-siRNA or FE65-siRNA for 24 h, respectively, were examined by immunoblot using specific antibodies. **(M-O)** Quantification of ULK1/GAPDH ratios (M), LC3II/LC3I ratios (N), P62/GAPDH ratios (O), respectively, (mean±SEM, *n*=3) (\*\**P*<0.01).

It has been well established that ATF4 is not only required for ER stress but also needed for hypoxia-induced expansion of autophagy, which is evidenced by the observation that ATF4 regulated the expression of LC3 ^28^ ^29^ ATG5 ^30^ and ULK1 ^31^. For this reason, we further examined the protein level of p62 and LC3 to determine whether overexpression of APP had any impact on autophagy. Immunoblot demonstrated that the protein level of ULK1 was decreased, while the protein level of p62 was increased in N2a SW cells compared with those in N2a cells. Meanwhile, the ratio of LC3II/LC3I was lower in N2a SW cells than that in N2a cells (Figure 4F-I), indicating that the autophagic flux was inhibited by APP overexpression. Immunofluorescence of LC3 also illustrated less conversion of LC3 in N2a SW cells relative to N2a cells (Figure 4J, K). However, when FE65 was knockdown by siRNA in both cell lines, unlike the recovery of ATF4, p62 and the ratio of LC3II/LC3I were not rescued, only the protein level of ULK1 was increased in N2A SW cells compared with N2a SW cells that were treated by scrambled control siRNA (Figure 4L-O), indicating that knockdown of FE65 was not sufficient to rescue the autophagy impairment induced by APP overexpression.

### 3.5 Knockout of APP facilitated LC3 conversion in H4 cells

Because the overexpressed APP695 contained a Swedish mutation, which might release more Aβ than wild-type APP, we could not simply draw the conclusion that the defects of autophagy in N2a SW cells were due to APP overexpression. In addition, FE65 knockdown by siRNA failed to rescue autophagy in the N2a SW cells. Thus, to verify the impact of APP on autophagy, we further established two H4 cell lines by CRISPR-Cas9, in which APP was ablated or introduced the Swedish mutation, respectively. Immunoblot verified the knockout of APP in H4 cells (Figure 5A, B). Though overexpression of APP upregulated the protein levels of FE65, GM130 and Erp72, the APP Swedish mutant and APP deficiency did not affect their protein levels in H4 cells (Figure 5A, C, D, E). Interestingly, the ratio of LC3II/LC3I was significantly increased in APP knockout H4 cells (H4-APPKO) compared with that in normal H4 cells (H4-control) and H4 cells harboring the APP Swedish mutant containing (H4-APPswe) (Figure 5A, G). However, the protein levels of ULK1 and p62 did not show significant differences (Figure 5A, F, H). Consistent with the immunoblot, immunofluorescence staining by using APP-CTF antibody showed that APP was located mainly on GA of normal H4 cells (Figure 5I, upper) and H4 cells harboring APP Swedish mutant (H4-APP swe) (Figure 5I, middle), but it was undetectable in APP KO cells (Figure 5I, lower). Of note, FE65 was not clustered in H4 cells **(Supplementary Figure2)**, instead, it was located at the protrusions of H4 cells. In addition, immunofluorescence of LC3 revealed that compared with normal H4 cells (Figure 5J, upper), the APP Swedish mutant did not impact the activation of LC3 (Figure 5J, middle), however, the conversion of LC3 was significantly increased in APP knockout H4 cells (Figure 5J, lower).

**Figure 5.**
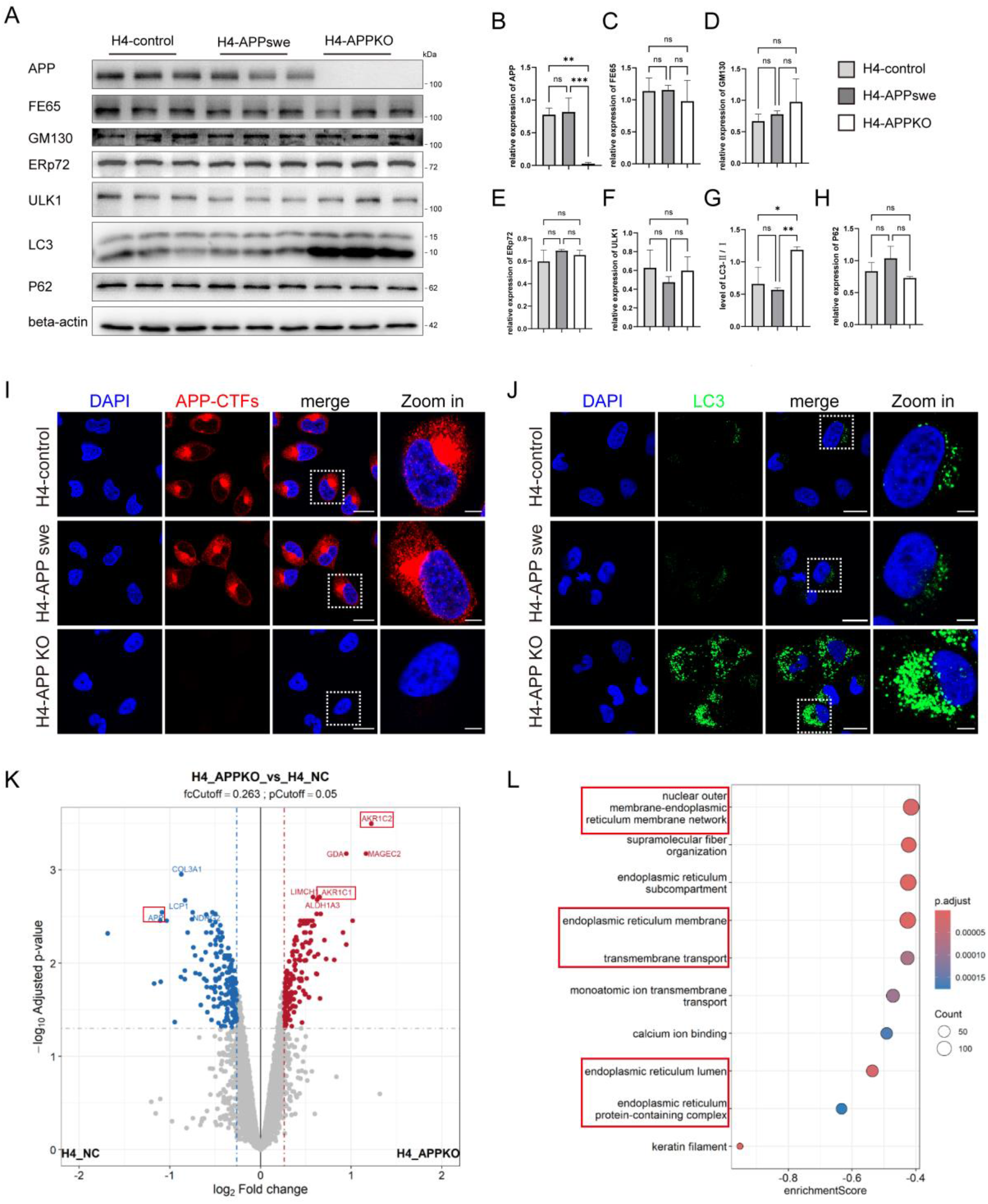
APP knockout facilitated LC3 conversion. **(A)** The protein levels of APP, FE65, GM130, Erp72, ULK1, LC3, and P62 in the lysates from normal H4 cells (H4 control), APP Swedish mutant containing H4 cells (H4-APPswe), and APP knockout H4 cells (H4-APPKO) were examined by immunoblot using specific antibodies. **(B-H)** Quantification of APP/actin ratios (B), FE65/actin ratios (C), GM130/actin ratios (D), Erp72/actin ratios (E), ULK1/actin ratios (F), LC3II/LC3I ratios (G), P62/actin ratios (H), respectively, (mean±SEM, *n*=3) (\**P*<0.05; \*\**P*<0.01; \*\*\*\**P*<0.0001). **(I)** Representative immunofluorescence staining using APP-CTF antibody (red) in normal H4 cells (H4 control, upper), APP Swedish mutant containing H4 cells (H4-APPswe, middle), and APP knockout H4 cells (H4-APPKO, lower). The nuclei were visualized by DAPI (blue). Scale bars, 20 μm in original or 5 μm in zoomed in. **(J)** Representative immunofluorescence staining using LC3 antibody (green) in normal H4 cells (H4 control, upper), APP Swedish mutant containing H4 cells (H4-APPswe, middle), and APP knockout H4 cells (H4-APPKO, lower). The nuclei were visualized by DAPI (blue). Scale bars, 20 μm in original or 5 μm in zoomed in. **(K)** Volcano plots of the proteome of APP knockout H4 cells (H4-APPKO) compared with that of normal H4 cells (H4-NC) (*n*=3, *P*<0.05, log^2^(Fold Change)>0.26). **(L)** Differentially expressed proteins between APP knockout H4 cells and normal H4 cells enriched in Gene Ontology (GO) terms with respect to their biological functions.

To further study the impact of APP ablation on H4 cells, we compared the proteome between APP knockout H4 cells and normal H4 cells. The results showed that AKR1C1, AKR1C2, et al., were upregulated in APP knockout cells, indicating that APP deficiency might impact steroid metabolism (Figure 5K, data are available at https://zenodo.org/records/23800849). Of note, we also found that overexpression of APP inhibited the protein level of SREBP2 **(Supplementary Figure 3A, B)**, which is in line with the report that APP controlled cholesterol turnover ^32^. GO analysis suggested that the differentially expressed proteins in APP knockout H4 cells were involved in regulating the reticulum membrane network and transmembrane transport (Figure 5L). These proteins were displayed in **supplementary Figure 3C**.

### 3.6 Swedish mutation facilitated cleavage of APP-CTF by γ-secretase

It has been demonstrated that the mutation of APP influenced its subcellular localization. The Swedish mutant APP was enriched in the GA compared with wild-type APP (APPwt). In contrast, APPice mutant (A673T in APP770), which was known to inhibit the generation of Aβ, was enriched in endosomes ^25^. Thus, we further detected the subcellular localization of APP by 4G8 antibody (Figure 6A, B), which recognizes the N-terminus of CTF-83 that is generated by the cleavage of α-secretase, and 6E10 antibody (Figure 6C, D), which recognizes the N-terminus of CTF-99 that is derived from the cleavage of β-secretase. The fluorescence of 4G8 was detected in the cytosol with puncta-like feature in normal H4 cells (H4 control, Figure 6A, upper). However, in those H4 cells that express APP Swedish mutant (H4-APP swe), very few of the fluorescence of 4G8 showed puncta-like feature. Instead, it became diminished (Figure 6A, lower). Consistent with 4G8, the fluorescence of 6E10 (Figure 6C, upper) was also detected in the cytosol of normal H4 cells as puncta, but presented as dot-like features in H4-APP swe (Figure 6C lower), which is similar to the feature of endosomes. These observations are similar with the previous reports that APPwt was preferentially to be transported from the Golgi apparatus to early endosomes ^25^. However, in this experiment, we predicted that both newly synthesized wild-type APP and APP Swedish mutant are located on GA (Figure 5J). Indeed, it is the CTF of wild-type APP that was favored to be transported to the lysosome compared with the CTF of APP Swedish mutant. These observations also suggested that newly synthesized APP did not undergo proteolysis until it had arrived at GA, which is consistent with the report that ER was probably not the proteolytic site for APP ^33^.

**Figure 6.**
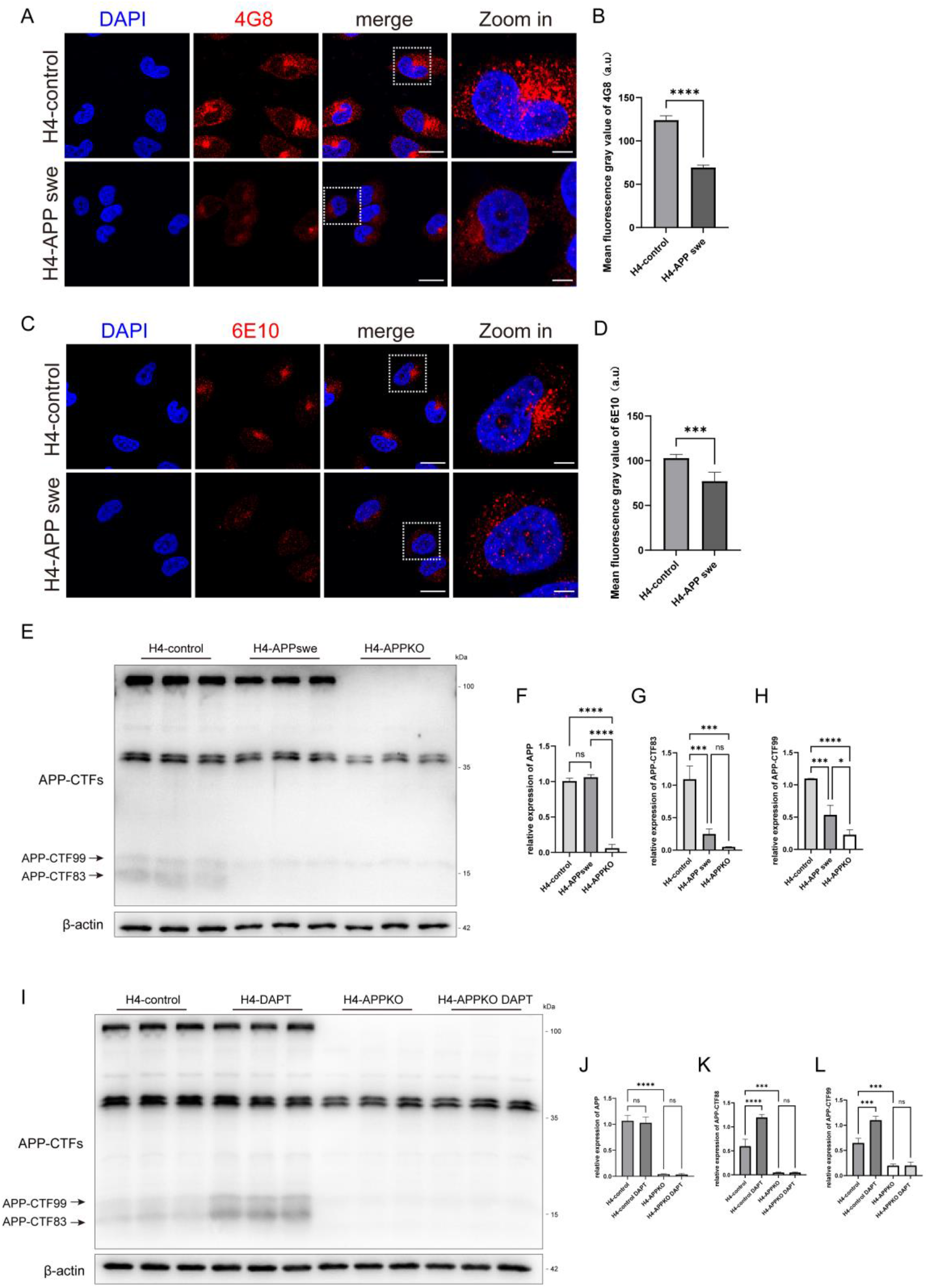
Swedish mutation of APP promoted CTF cleavage by γ-secretase. **(A)** Representative immunofluorescence staining using 4G8 antibody (red) in normal H4 cells (H4 control, upper), APP Swedish mutant containing H4 cells (H4-APPswe, lower). The nuclei were visualized by DAPI (blue). Scale bars, 20 μm in original or 5 μm in zoomed in. **(B)** Comparison of the fluorescence intensity of 4G8 between normal H4 cells (H4 control) and APP Swedish mutant containing H4 cells (H4-APPswe). (mean±SEM, *n*=8) (\*\*\*\**P*<0.0001). **(C)** Representative immunofluorescence staining using 6E10 antibody (red) in normal H4 cells (H4 control, upper), APP Swedish mutant containing H4 cells (H4-APPswe, lower). The nuclei were visualized by DAPI (blue). Scale bars, 20 μm in original or 5 μm in zoomed in. **(D)** Comparison of the fluorescence intensity of 6E10 between normal H4 cells (H4 control) and APP Swedish mutant containing H4 cells (H4-APPswe). (mean±SEM, *n*=8) (\*\*\**P*<0.001). **(E)** The protein levels of APP and APP-CTFs in the lysates from normal H4 cells (H4-control), APP Swedish mutant containing H4 cells (H4-APPswe), and APP knockout H4 cells (H4-APPKO) were examined by immunoblot using APP-CTF antibodies. **(F-H)** Quantification of APP/actin ratios (F), CTF-83/actin ratios (G), CTF-99/actin ratios (H), (mean±SEM, *n*=3) (\**P*<0.05; \*\*\**P*<0.001; \*\*\*\**P*<0.0001). **(I)** The protein levels of APP-CTFs in the lysates from normal H4 cells (H4-control) APP knockout H4 cells (H4-APPKO) that were treated with DMSO or DAPT in equal amount of DMSO (final concentration of DAPT 100 nM), respectively, were examined by immunoblot using APP-CTFs antibodies. **(J-L)** Quantification of APP/actin ratios (J), CTF-83/actin ratios (K), CTF-99/actin ratios (L), (mean±SEM, *n*=3) (\*\*\**P*<0.001; \*\*\*\**P*<0.0001).

The intracellular trafficking and proteolytic processing of APP were quite controversial. Tam et al., ^34^ and Lorenzen et al.,^35^ suggested that lysosome is a likely site of Aβ production, and the Swedish mutation caused rapid clearance of APP and generation of Aβ from lysosomes. However, Choy et al., observed that Aβ is generated predominantly in trans-Golgi network (TGN), which relies on the recycling of APP from early endosomes to the TGN ^36^ in HEK293 cells. Xia et al., ^37^ and Thinakaran et al., ^38^ also provided strong evidences for that APP is processed to generate Aβ at the TGN. In addition, Tan et al., identified that TGN is the major site for α-secretase processing of APP in HeLa cells and primary neurons as well ^39^. Moreover, Toh, et al. suggested that the newly synthesized APP is directly transport from the TGN to the early endosomes for degradation ^40^. To illustrate the influence of Swedish mutation on APP proteolytic process, we further compared the protein levels of APP-CTFs by immunoblot, and consistent with the observations by immunofluorescence, the blots showed that compared with wild-type APP, APP Swedish mutant showed similar level of full-length APP (Figure 6E, F), but produced smaller amounts of CTF-99 and CTF-83 (Figure 6E, G, H). The clearance of CTF by γ-secretase was further demonstrated by adding DAPT, the inhibitor of γ-secretase, in both H4 cells and APP knockout H4 cells, respectively. DAPT significantly increased the protein level of APP-CTF in H4 cells but did not impact the protein level of full-length APP (Figure 6I-L). This result suggested that APP Swedish mutant promoted the cleavage of APP-CTF by γ-secretase.

## 4 Discussion

The mutations of *APP* are closely correlated with the development of AD. Since the Amyloid cascade hypothesis was proposed in 1992 ^41^, many efforts have been made to illustrate the neurotoxicity of Aβ and tau aggregates. However, the underlying mechanism of how Aβ induced tau phosphorylation and aggregation is still unknown, as well as how do they trigger neurocyte death today. We believe that demonstrating the biological function of APP and tau may contribute to understanding the etiology of AD.

In the current study, we found that overexpression of APP695 resulted in the retention of APP in GA and ER, and promoted FE65 clustering outside the nuclei. Moreover, the quantitative change of APP caused qualitative changes of FE65 in regulating ATF4 transcription. These observations are consistent with previous studies that manifested that APP colocalize with FE65 in the ER/GA and possibly in endosomes ^42^, and served as a cytosolic anchoring site that prevents FE65 nuclear translocation ^43^. Sabine Probst et al., proposed that the rate of APP cleavage regulates the nuclear translocation of AICD–Fe65–Tip60 (AFT) complexes to promote the transcriptional activity of Fe65 ^44^. Interestingly, we observed that by promoting FE65 clustering, APP overexpression prevented ER stress which is evidenced by the downregulated protein level of ATF4 and its downstream effectors, including Bcl-2 and CHOP. More importantly, knockdown of FE65 rescued the protein level of ATF4, as well as Bcl2 and Chop. These results argued that the clustering of FE65 outside the nuclei also affected gene transcription, the underlying mechanism might be that it indirectly segregation of other transcription factors from entering the nucleus. Interestingly, the mRNA level of APP was lowest in newborn rat brain, whereas reached the highest at P20-P40 ^45,46^. It is also of interest to learn whether APP promotes FE65 clustering during neuronal development in future studies.

On the other hand, the overexpression of APP also impaired autophagy, besides stabilizing GA and ER, which is not dependent on FE65 in our experiments. This phenomenon was also in agreement with the dysfunction of lysosome found in AD model mice and Down syndrome fibroblasts ^47–49^. In contrast, knockout of APP remarkably facilitated LC3 conversion. These observations are also in line with the report that deficiency of APP rescued the lysosome defects induced by PSEN1 mutations ^50^. Of note, besides brain, APP was also highly expressed in islets ^51^. Overexpression of APP was found to stimulate insulin secretion ^51^, on the contrary, APP knockout mice showed decreased insulin level ^52^ ^53^. Therefore, whether APP regulated insulin content through impacting autophagy needs to be further studied.

APP overexpressing mice models were widely used to mimic the pathology of AD. In the current study, however, we learned that except for producing large amounts of Aβ, overexpression of APP could also promote FE65 clustering outside the nuclei. These observations suggested that the quantitative changes of APP played an important role in endomembrane system during development. Therefore, the APP mutant knock-in models, instead of APP overexpression model, might be more suitable for the Aβ neurotoxicity related studies. In addition, in this study we also noticed that the APP Swedish mutant significantly promoted the proteolysis of APP-CTF by γ-secretase, which accompanied by less CTF transported to lysosome. Previous studies demonstrated that overexpression of APP Swedish mutant significantly increased the level of APP-CTF99 ^38,54^. However, according to the observations in the current study, we proposed an “APP-CTFs escape hypothesis”, we predicted that at normal conditions, CTF-99 and CTF-83 of APP were mainly degraded by lysosome as evidenced by autophagy inhibition induced APP-CTFs accumulation ^55^, only a small portion of Aβ needs to be eliminated by IDE and NEP ^56^. However, the APP Swedish mutant not only increased the β-cleavage of APP but also triggered abnormal cleavage of CTFs by γ-secretase which resulted in the splitting of Aβ fragment from AICD. Of note, the AICD contains a KFERQ motif, which is responsible for the chaperone-mediated autophagy (CMA) mediated lysosome-targeting degradation of APP ^57^. Thus, the dissociation of Aβ and AICD could induce undesirable escape of Aβ from lysosome degradation, in turn give rise to Aβ accumulation. This hypothesis also requires further verification in future work.

As mentioned above, revealing the biological function of APP and tau may contribute to not only understanding the etiology of AD, but also providing therapeutic approaches for the related diseases. Previously, we reported that human tau promoted Warburg effect-like glycolytic metabolism under acute high glucose conditions ^58^. This study suggested that efforts should focus on prohibiting tau aggregation rather than eliminating tau. Because tau plays an important role in maintaining homeostasis of NAD^+^ and ROS. In this study, we demonstrated that overexpression of APP not only contributed to stabilizing GA and ER, but also induced stagnation of autophagy and impaired cholesterol synthesis, which is also supported by previous studies showing that APP is involved in regulating lipoprotein metabolism ^59^. However, since knockout of APP might facilitate autophagy which will contribute to alleviating tau accumulation, eliminating APP by siRNA or ASO may be a feasible method for preventing the development of AD.

## Acknowledgments

We thank the Instrumental Analysis Center of Shenzhen University (Lihu Campus) for offering access to the laser scanning confocal microscopes (LSM 710 and LSM 880, ZEISS, Germany). We are also grateful for the support from the public service platform of instruments and equipment at the College of Life Sciences and Oceanography of Shenzhen University.

## Conflicts of Interest

All the authors listed in the manuscript declare that no conflict of interest exists.

## Funding Sources

National Natural Science Foundation of China (31700919, 82674996); Guangdong Provincial Natural Science Foundation (2025A1515011129), Guangdong Basic and Applied Basic Research Foundation (2026A1515010558), Shenzhen-Hong Kong Institute of Brain Science-Shenzhen Fundamental Research Institutions (2023SHIBS0003).

## Consent Statement

Not applicable for this study. Consent was not necessary.

## Notes

### Competing Interest Statement

The authors have declared no competing interest.

