## Supplementary material for "APP functions at the endomembrane system": Tables

Table 1. Antibodies used in this study.

| Antibodies | Cat No. | Supplier | Species |
| --- | --- | --- | --- |
| ULK1 | 8054S | Cell Signaling Technology | Rat |
| LC3 | M18P1 | Selleck Chemicals | Rat |
| P62 | 31403-1-AP | Proteintech | Rat |
| GM130 | 11308-1-AP | Proteintech | Rat |
| Erp72 | 14712-1-AP | Proteintech | Rat |
| FE65 | 2877S | Cell Signaling Technology | Rat |
| APP-CTFs | A8717 | Merck | Rat |
| 6E10 | MA5-51794 | ThermoFisher Scientific | Mouse |
| 4G8 | MAB1561-M | Merck | Mouse |
| SREBP2 | 28212-1-AP | Proteintech | Rat |
| ATF4 | D4B8 | Cell Signaling Technology | Rat |
| CHOP | L63F7 | Cell Signaling Technology | Rat |
| eIF2a | #9722 | Cell Signaling Technology | Rat |
| p-eIF2a | #9721 | Cell Signaling Technology | Rat |
| Bcl-1 | 3498S | Cell Signaling Technology | Rat |
| Beta-actin | 66009-1-lg | Proteintech | Mouse |
| GAPDH | 10494-1-AP | Proteintech | Rat |

Table 2. The spacer sequence of sgAPP.

| Name | Sequence (5’→3’) |
| --- | --- |
| spacer of sgAPP | GGAGAUCUCUGAAGUGAAGA |
| ssODN donor | ATACTTTAATTATGATGTAATACAGGTTCTGGGTTGACAAATATCAAGACGGAGGAGATCTCTGAAGTGAATCTGGATGCAGAATTCCGACATGACTCAGGATATGAAGTTCATCATCAAAAATTGGTACGTAAAATAATTTA |

Table 3. Primers used in amplifying Cas9 cleavage site in genomic region.

| Gene | Sequence (5’→3’) |
| --- | --- |
| APP | F: ATAAGCCCGTAAGCCAAGCC  R: TTCAAGCAAGTCTCCTGCGT |

F: forward primer; R: reverse primer.
