## supplementary figures for "APP functions at the endomembrane system"

Figure.S1

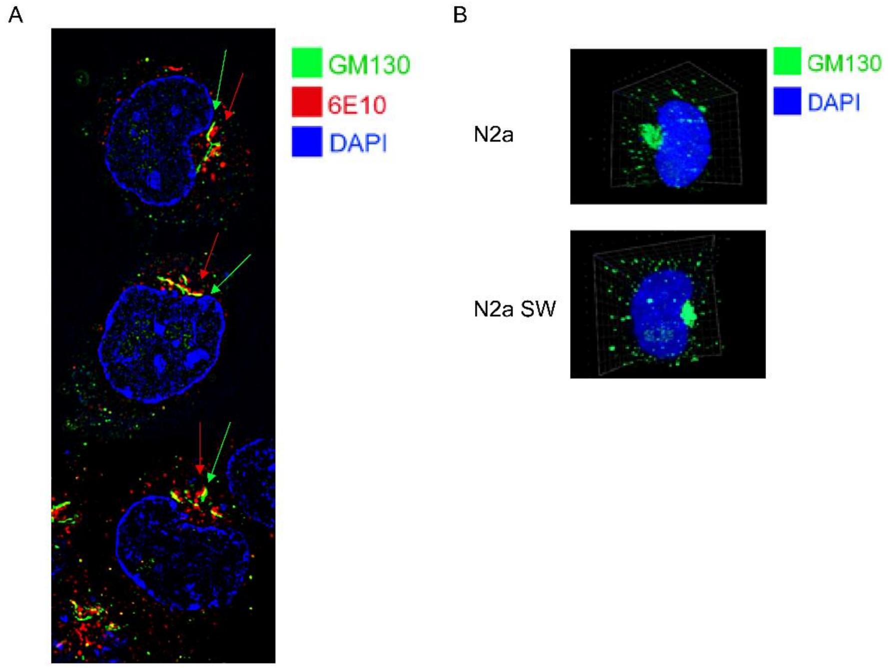


**Supplementary Figure 1. Overexpressed APP were localized on trans-Golgi (A)** Representative image of the immunofluorescence staining using GM130 (green) and 6E10 (red) antibodies in normal N2a SW cells that captured by Structured Illumination Microscopy (SIM). The nuclei were visualized by DAPI (blue). **(B)** Representative immunofluorescence staining using GM130 antibody (green) in N2A cells (upper) and N2a SW cells (lower). The nuclei were visualized by DAPI (blue)**.**

Figure.S2

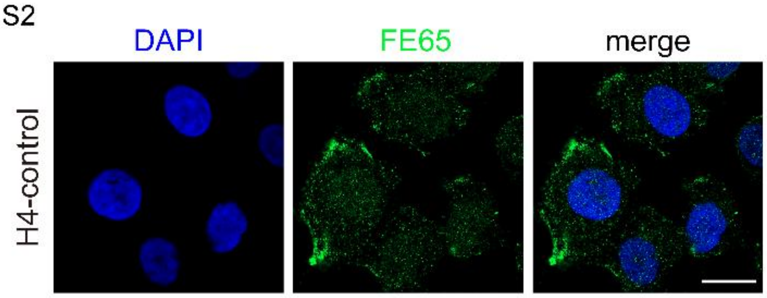


**Supplementary Figure 2. Subcellular localization of FE65 in H4 cells.** Representative immunofluorescence staining using fe65 (green) in normal H4 cells. The nuclei were visualized by DAPI (blue).

Figure.S3

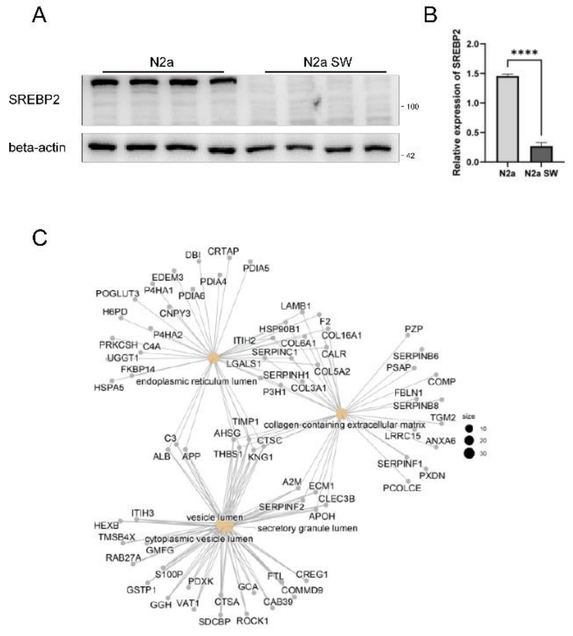


**Supplementary Figure 3. APP overexpression and knockout influenced ER lumen associated proteins (A)** The protein levels of SREBP2 in the lysates from normal N2a cells and N2a SW cells were examined by immunoblot using SREBP2 specific antibodies. **(B)** Quantification of SREBP2/actin ratios. (mean±SEM, *n*=4) (*****P*<0.0001). **(C)** The differently expressed protein in APP knockout H4 cells compared with normal H4 cells were analyzed by Gene Ontology (GO) enrichment approach. Those were related with endoplasmic reticulum lumen, vesicle lumen, cytoplasmic vesicle lumen, and collagen-containing extracellular matrix were displayed in the graph.
